# Astrocytic TIMP-1 regulates production of Anastellin, a novel inhibitor of oligodendrocyte differentiation and FTY720 responses

**DOI:** 10.1101/2023.02.17.529003

**Authors:** Pearl A. Sutter, Cory M. Willis, Antoine Menoret, Alexandra M. Nicaise, Anthony Sacino, Arend. H. Sikkema, Evan Jellison, Kyaw K. Win, David K. Han, William Church, Wia Baron, Anthony T. Vella, Stephen J. Crocker

## Abstract

Astrocyte activation is associated with neuropathology and the production of tissue inhibitor of metalloproteinase-1 (TIMP1). TIMP1 is a pleiotropic extracellular protein that functions both as a protease inhibitor and as a growth factor. We have previously demonstrated that murine astrocytes that lack expression of *Timp1* do not support rat oligodendrocyte progenitor cell (rOPC) differentiation, and adult global *Timp1* knockout (*Timp1*^KO^) mice do not efficiently remyelinate following a demyelinating injury. To better understand the basis of this, we performed unbiased proteomic analyses and identified a fibronectin-derived peptide called anastellin that is unique to the murine *Timp1*^KO^ astrocyte secretome. Anastellin was found to block rOPC differentiation *in vitro* and enhanced the inhibitory influence of fibronectin on rOPC differentiation. Anastellin is known to act upon the sphingosine-1-phosphate receptor 1 (S1PR1), and we determined that anastellin also blocked the pro-myelinating effect of FTY720 (or fingolimod) on rOPC differentiation *in vitro*. Further, administration of FTY720 to wild-type C57BL/6 mice during MOG_35-55_-EAE ameliorated clinical disability while FTY720 administered to mice lacking expression of *Timp1* in astrocytes (*Timp1*^cKO^) had no effect. Analysis of human *TIMP1* and fibronectin (*FN1*) transcripts from healthy and multiple sclerosis (MS) patient brain samples revealed an inverse relationship where lower *TIMP1* expression was coincident with elevated *FN1* in MS astrocytes. Lastly, we analyzed proteomic databases of MS samples and identified anastellin peptides to be more abundant in the cerebrospinal fluid (CSF) of human MS patients with high versus low disease activity. The prospective role for anastellin generation in association with myelin lesions as a consequence of a lack of astrocytic TIMP-1 production could influence both the efficacy of fingolimod responses and the innate remyelination potential of the the MS brain.

**Significance Statement:** Astrocytic production of TIMP-1 prevents the protein catabolism of fibronectin. In the absence of TIMP-1, fibronectin is further digested leading to a higher abundance of anastellin peptides that can bind to sphingosine-1-phosphate receptor 1. The binding of anastellin with the sphingosine-1-phosphate receptor 1 impairs the differentiation of oligodendrocytes progenitor cells into myelinating oligodendrocytes *in vitro*, and negates the astrocyte-mediated therapeutic effects of FTY720 in the EAE model of chronic CNS inflammation. These data indicate that TIMP-1 production by astrocytes is important in coordinating astrocytic functions during inflammation. In the absence of astrocyte produced TIMP-1, elevated expression of anastellin may represent a prospective biomarker for FTY720 therapeutic responsiveness.

## Introduction

Reactive astrocytes are common to all neurological diseases and possess a distinct transcriptome from astrocytes in the healthy CNS (1, 2). The influence of astrocytes on the pathology of the autoimmune demyelinating disease multiple sclerosis (MS) is being increasingly recognized as a central component of the disease process, with dual roles as both mediators of the neuroimmune response (3–6) and as a source of factors with the potential to either promote or impair myelin regeneration (7–11). One such factor is tissue inhibitor of metalloproteinase (TIMP)-1, which has been shown to be produced by both human and mouse reactive astrocytes (12).

Tissue inhibitor of metalloproteinase (TIMP)-1 is a secreted extracellular protein with dual functions as a protease inhibitor and as a growth factor (13, 14). Expression of TIMP-1 in the developing embryonic and early postnatal mouse and rat central nervous systems (CNS) is higher than in the healthy adult CNS (15–17), where TIMP-1 is expressed at very low to undetectable levels (15, 17). However, injury or infection in the adult CNS rapidly and robustly induces TIMP-1 expression by astrocytes (15, 18, 19). In experimental murine models and in humans with MS, astrocytes expressing TIMP-1 are localized to the periphery of demyelinating lesions (15, 20, 21). Interestingly, mice with global *Timp1* gene knockout (*Timp1*^KO^)exhibit delayed CNS myelination during development (22). Adult *Timp1*^KO^ mice also exhibit chronically demyelinated lesions in the experimental autoimmune encephalomyelitis (EAE) mouse model of CNS inflammation (23, 24) and cuprizone-induced demyelination (21). Conversely, transgenic mice that over-express TIMP-1 in astrocytes have reduced white matter injury in EAE (25). Thus, the temporal and spatial expression of elevated TIMP-1 by astrocytes during inflammation-induced demyelination is associated with maintaining CNS myelin integrity while its absence is associated with chronic demyelination (14).

It is recognized that the environment of the demyelinated lesion is also a source of factors which can directly limit the remyelinating capacity of the CNS in diseases like MS. The cellular environment of the demyelinated lesion is also variable, which can contain lymphocytes and active phagocytes, but reactive astrocytes are a consistent feature. Importantly, peri-lesion astrocytes abundantly produce the extracellular matrix (ECM) molecule fibronectin (Fn) (26). Fn is a large glycoprotein that plays an important role in the ECM due to its ability to bind other ECM molecules, such as collagen and proteoglycans, as well as other Fn molecules; the binding of Fn to other Fn molecules results in the formation of a structure referred to as the “fibrillar matrix”(27, 28). Functions of Fn include cell adhesion, migration, and differentiation (27). Fn produced by astrocytes during demyelinating stages of MS (26), can form high molecular weight aggregates which are postulated to be the result of inefficient clearing following injury. In EAE, Fn can sustain demyelination by inhibiting the differentiation of oligodendrocyte progenitor cells (OPCs) to myelinating oligodendrocytes (26, 29). Whether astrocytic TIMP-1 expression affects the generation of Fn has not been previously studied.

Herein, we have tested how astrocyte TIMP-1 affects OPC maturation, using both *in vitro* and *in vivo* models, by studying a novel *Gfap-CRE/Timp1^fl/fl^* mouse line which lacks astrocytic expression of TIMP-1 (*Timp1*^cKO^). A proteomic analysis of the astrocyte secretome from wildtype and *Timp1*^cKO^ primary astrocytes identified increased abundance of the Fn and an Fn-derived peptide called Anastellin. Rat OPCs treated with recombinant anastellin led to a significant reduction in their maturation into myelin basic protein (MBP) expressing mature oligodendrocytes. This reduction was observed even in the presence of the pro-myelinating drug FTY720. However, addition of recombinant TIMP-1 to the culture media removed this block and restored the maturation of OPCs to oligodendrocytes. *Timp1*^cKO^ mice with MOG_35-55_ EAE had a significantly worse clinical disease course that was refractory to FTY720 treatment. Analysis of human brain tissue and cerebrospinal spinal fluid (CSF) from control and patients with MS revealed an increased fibronectin to TIMP1 protein ratio in MS patients and increased abundance of the peptide anastellin, respectively. Together, these results implicate expression of TIMP-1 in the regulation of astrocytic Fn with implications for oligodendrocyte biology.

## Results

### Astrocytic TIMP-1 knockout mice develop enhanced EAE phenotype

To address the contribution of astrocytic TIMP-1 to the clinical course of MOG_35-55_-EAE we developed a novel conditional *Timp1* deficient mouse (*Timp1*^cKO^)(**Fig. 1*A***) in which *Timp1* expression is specifically deleted from astrocytes under the control of the *Gfap* promoter (*Gfap-* CRE/*Timp1*^fl/fl^) (30). Using primary glial cultures from P0-P3 *Timp1*^cKO^ mice, we verified CRE expression in GFAP+ astrocytes using immunocytochemistry (**Fig. 1*B***). To verify the functional loss of *Timp1* expression from *Timp1*^cKO^ astrocytes, wild-type (WT) and *Timp1*^cKO^ primary glia were challenged with interleukin-1β (IL-1β, 10 ng/ml), a potent inducer of *Timp1* expression in astrocytes (31). Consistent with previous studies, IL-1β treatment led to a significant induction of *Timp1* mRNA expression in WT astrocytes whereas IL-1β had no effect on *Timp1*^cKO^ astrocytes (**Fig. 1*C***). These data confirmed the efficacy of the cell-specific gene deletion *in vitro*.

**Figure 1.**
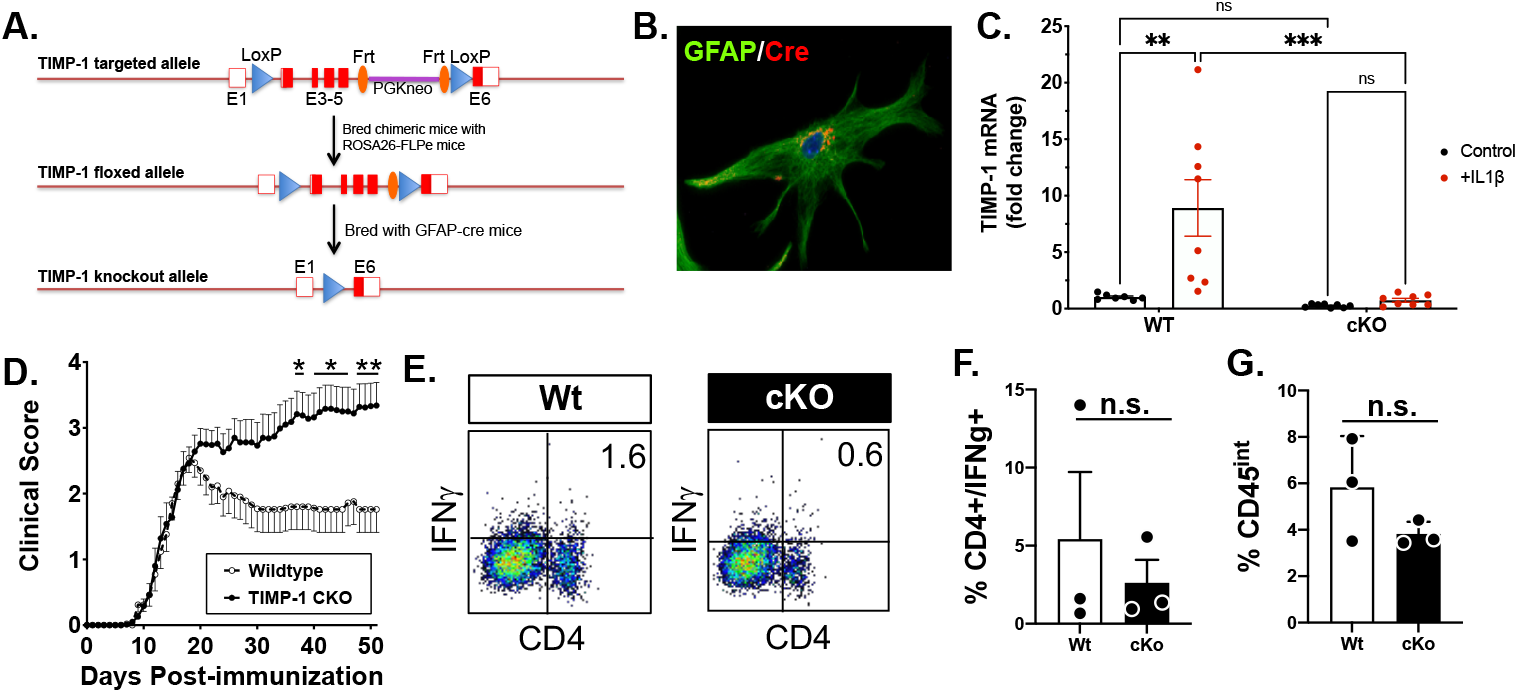
Astrocyte-specific TIMP-1 knockout mice develop a severe EAE phenotype that is immunologically independent. **(A)** Schematic of design of development of the flox-targeted sites that enabled cell and tissue-specific deletion of TIMP-1 in GFAP-Cre-TIMP-1^fl/fl^ mice. **(B)** Immunocytochemical validation of CRE expression in astrocytes from GFAP-Cre:TIMP-1^fl/fl^ mice, and **(C)** physiological validation of TIMP-1 mRNA expression in primary astrocytes and absence of TIMP-1 mRNA expression in astrocyte cultures from GFAP-Cre:TIMP-1^fl/fl^ mice (ANOVA; ****, P<0.0001; where *,P<0.01; ***, P<0.002). **(D)** Clinical EAE in MOG_35-55_-treated C57Bl/6 wildtype and GFAP-Cre:TIMP-1^fl/fl^ mice over 52 day time course. **(E,F)** Flow cytometry analysis of overall CD4+/IFNγ+ T cells from spinal cords of Wt and GFAP-Cre:TIMP-1^fl/fl^ mice mice at peak clinical EAE illness (day 18), including **(F)** analysis of CD4+/IFNγ+ T cells (t-test, P<0.20) and **(G)** CD45^int^/CD11b+ microglia (t-test, P<0.285) in each genotype.

We then wanted to assess the functional contribution of astrocytic TIMP-1 in the MOG_35-55_-EAE mouse model of inflammatory demyelination (23). *Timp1*^cKO^ and WT littermates were immunized with MOG_35-55_ peptide to induce EAE, and the development of clinical disease was assessed daily. *Timp1*^cKO^ and WT mice did not differ in time of clinical onset or peak clinical severity (~18 days post immunization, dpi), however *Timp1*^cKO^ mice developed a persistent, severe clinical course of EAE that was sustained throughout the study period (52 days) (**Fig. 1*D***). Clinical disease in this MOG_35-55_ EAE model is predominantly driven by CD4+ T cells (32, 33). Therefore, to understand if the increased disease severity is due to an expansion and persistence of this cell population during the chronic stage of EAE (day 50) we evaluated the magnitude of CD4+/IFNγ+ T cells in the CNS by flow cytometry. We found no significant differences in the proportion of CD4+/IFNγ+ T cells between WT and *Timp1*^cKO^ mice (**Fig 1*E*, 1*F***), which is consistent with our previous study that knockout of TIMP-1 does not influence the peripheral immune response in EAE mice (23). In contrast, the increased response of CD11b+/CD45^in^ microglia, which we previously reported on in *Timp1*^KO^ mice with EAE (23), was not apparent in *Timp1*^cKO^ mice as assessed by flow cytometry (**Fig 1*G***).

### Proteomic Analysis of the *Timp1*^cKO^ Astrocyte Secretome Identifies Dysregulation of Fibronectin

We previously determined that conditioned media from *Timp1*^KO^ primary glial cultures did not support the differentiation and maturation of rat oligodendrocyte progenitor cells (rOPCs) into mature MBP+ oligodendrocytes *in vitro(22*). Therefore, we hypothesized that the sustained clinical disability observed in the *Timp1*^cKO^ EAE mice was a result of an altered astrocyte secretome that prevented the regeneration of the demyelinated spinal cord. To address this question, astrocyte conditioned media (ACM) from WT and *Timp1*^cKO^ primary glia cultures was collected and analyzed by an automated 2D protein fractionation system (34). Post-separation analysis of protein abundance peaks using proteome lab software revealed unique differences in the secretome of WT and *Timp1*^cKO^ astrocytes (**Fig. 2*A*, 2*B***). Mass spectrometry analysis of six unique peaks from the WT and *Timp1*^cKO^ ACM were used to compare time constant matched samples to identify a list of potential candidates (**Fig. 2*C*, 2*D***). Among the uniquely expressed peptides identified were fragments of fibronectin (Fn).

**Figure 2.**
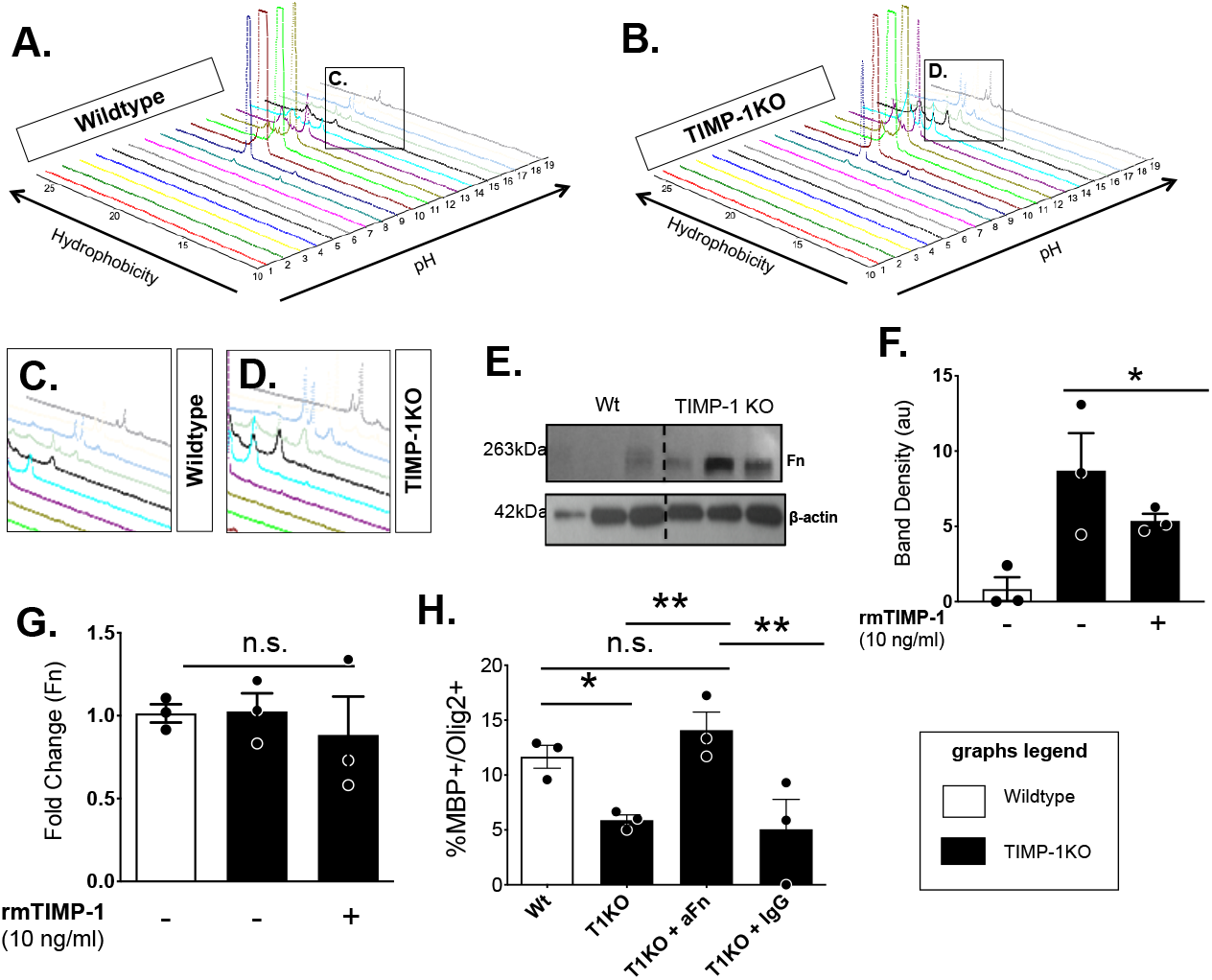
Proteomic identification of elevated fibronectin (Fn) in secretome of TIMP-1KO astrocytes. **(A,B)** Topographic projections rendered using ProteomeLab™ of PF-2D analyses of extracellular proteins in secretome of primary astrocytes from wildtype (A,C) and TIMP-1KO (B,D) cultures. **(C,D)** Magnified view of differential peaks of interest identified by ProteomeLab™ which were interrogated by LC-MS/MS. **(E)** Western blot analysis of fibronectin (Fn) protein expression in astrocyte conditioned media CM from wildtype (Wt) and TIMP-1KO astrocytes. **(F)** Quantification of Fibronectin in Wt, TIMP-1KO and TIMP-1KO cultures after treatment with recombinant murine TIMP-1. **(G)** Analysis of Fn mRNA expression by qPCR demonstrated no TIMP-1 regulation of Fn mRNA expression in primary TIMP-1KO astrocyte cultures. **(H)** Effect of immunodepletion of Fn from astrocyte conditioned media on OL differentiation in vitro in TIMP-1KO and Wt (control) treated cultures. Data in F,G,H were analyzed by ANOVA with either Bonferroni or Fisher’s LSD post-hoc tests where appropriate *, P<0.05, **, P<0.008 and ***, P<0.0005. Graphs represent data from n=3-6 technical replicates from an experiment which was representative of n=3 biologically replicated experiments.

Astrocytic Fn has been implicated as an impediment to remyelination in MS, implicating a potential inverse relationship between astrocytic TIMP-1 and Fn. To verify that Fn protein increased in *Timp1*^cKO^ astrocytes, we performed an immunoblot on cell lysates from WT and *Timp1*^cKO^ primary astrocyte cultures and confirmed elevated levels of Fn in *Timp1*^cKO^ vs WT astrocytes (**Fig. 2*E***). We then tested whether the increased abundance of Fn in the *Timp1*^cKO^ astrocytes is regulated in a TIMP1-dependent manner. Exogenous application of recombinant murine TIMP1 (rmTIMP1, 10 ng/mL) led to a significant reduction in the abundance of Fn in *Timp1*^cKO^ ACM (**Fig. 2*F***). The reduction in secreted Fn was not a result of transcriptional or post-transcriptional regulation by the exogenously added rmTIMP1 as the expression of *Fn1* mRNA in both WT and *Timp1*^cKO^ was not significantly different (**Fig. 2*G***). These data suggest that TIMP1 does not transcriptionally regulate *Fn1*, rather the accumulation of extracellular Fn peptides in the ACM results from an absence of TIMP1.

### Astrocyte produced fibronectin impairs the differentiation of OPCs into MBP+ oligodendrocytes

To define a functional role for the increased extracellular Fn from *Timp1*^cKO^ astrocytes, we tested whether immunodepletion of Fn in ACM using polyclonal antisera against Fn or IgM2 control antisera affected OPC differentiation. Consistent with our previous findings (11), *Timp1*^cKO^ ACM and *Timp1*^cKO^ ACM pre-treated with control IgM2 blocked the differentiation of rOPCs. However, immunodepletion of Fn from the *Timp1*^cKO^ ACM restored rOPC differentiation to comparable levels observed in rOPCs grown in WT ACM (**Fig. 2*H***). These data suggest that the increased extracellular abundance of Fn in *Timp1*^cKO^ ACM impairs the differentiation of rOPCs.

### Fibronectin breakdown peptide Anastellin blocks OPC differentiation

Among the Fn peptides identified in our proteomics screen of the *Timp1*^cKO^ ACM was a particularly abundant peptide called Anastellin (UniportKB-P11276). Anastellin is a proteolytic product derived from the C-terminus of the first type III domain of Fn (35). Anastellin is present in the blood and has known functions as an antagonist of angiogenesis and for facilitating the formation of high molecular weight (HMW) aggregates of Fn in demyelinated lesions in MS (26, 36). This indicates a possible connection between Anastellin and the inhibitory effects of Fn on rOPC differentiation. Exogenous application of recombinant murine fibronectin (rmFn; 1 and 10 μg/mL) to rOPCs impaired their differentiation in a concentration dependent manner compared with untreated controls (**Fig 3*A*, 3*B***). We then tested whether Anastellin alone or in combination with Fn could synergistically impair rOPC differentiation. We found that increasing concentrations of rmFn in combination with a consistent concentration of Anastellin resulted in a concentration dependent decrease in OL differentiation (**Fig 3*C*, 3*D***). Similarly, co-treatment of rOPCs with a concentration of rmFn that did not have a pronounced effect on impairing differentiation (1μg/ml) and increasing concentrations of rmAna, resulted in a concentrationdependent reduction in OL differentiation (**Fig. 3*C*, 3*E***). These data suggest that Anastellin enhances the effect of Fn on OPC differentiation.

**Figure 3.**
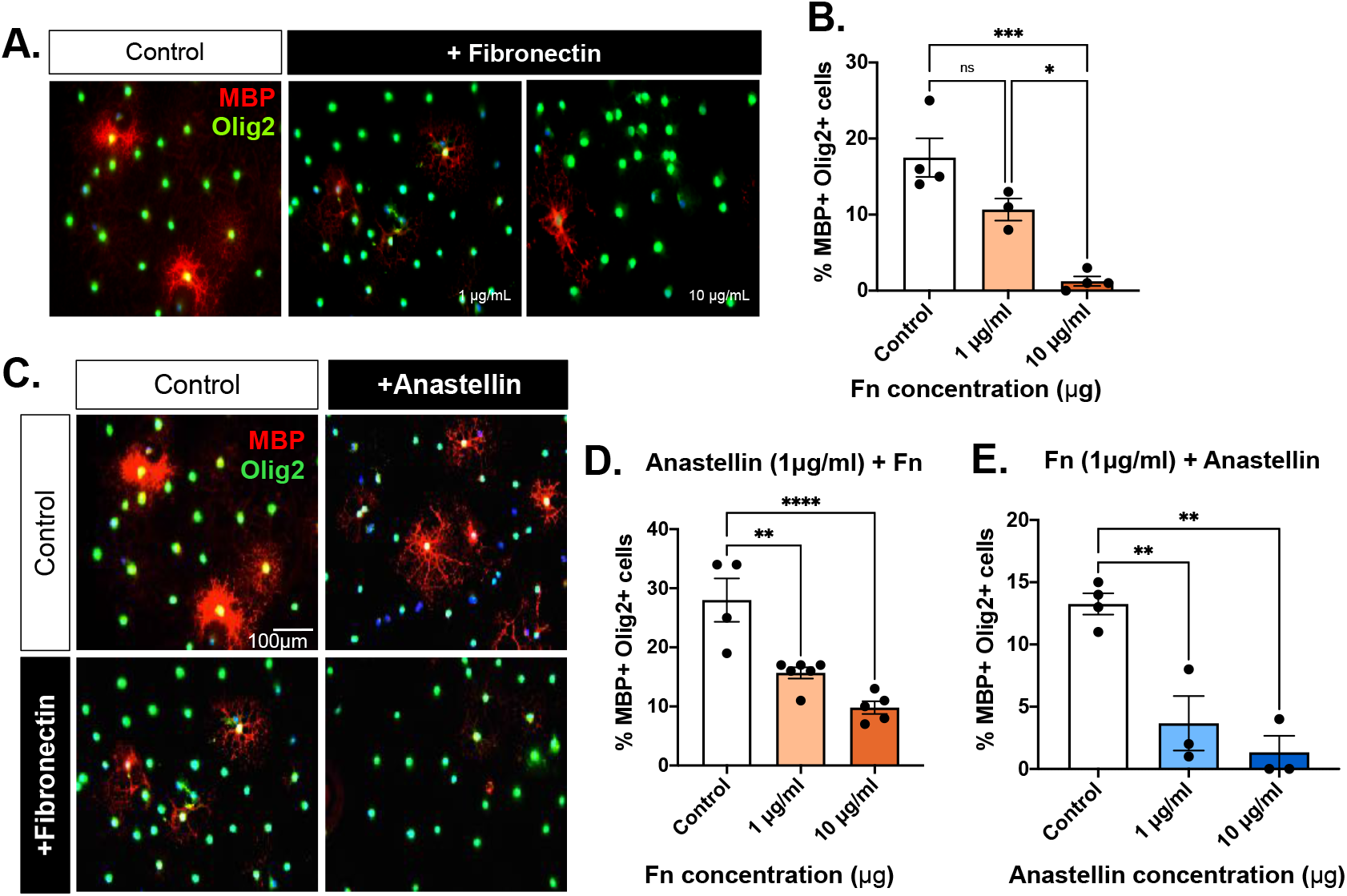
Anastellin enhances the inhibitory effect of Fibronectin on oligodendrocyte differentiation and blocks enhanced differentiation in response to FTY720. **(A)** Immunocytochemistry for mature oligodendrocytes (MBP+/Olig2+) under control differentiating conditions and with increasing concentrations of rmFibronectin (1 μg/ml, 10 μg/ml). **(B)** Quantification of OL differentiation following treatment with increasing concentrations of rmFibronectin. **(C)** Immunocytochemistry for mature oligodendrocytes (MBP+/Olig2+) under control differentiating conditions and with treatment of rmAnastellin (μg/ml), rmFibronectin (μg/ml) or both rmAnastellin and rmFibronectin. **(D)** Analysis of OL differentiation in primary OPC cultures treated with a constant concentration of rmFn (1 μg/m) but with increasing concentrations of rmAnastellin (1 μg/ml, 10 μg/ml), and **(E)** Analysis of OL differentiation in primary OPC cultures treated with constant concentration of rmAnastellin (1 μg/ml) but with increasing concentrations of rmFn (1 μg/ml, 10 μg/ml). Data in B (P<0.001), C (P<0.001) were analyzed by ANOVA with multiple-comparisons post-hoc tests where **, P<0.001 and ****, P<0.0001. Data in F was analyzed by t-test relative to baseline differentiation to determine significance **, P<0.008. Graphs represent data from n=3-6 technical replicates from an experiment which was representative of n=3 biologically replicated experiments.

### *Timp1*^cKO^ EAE mice do not respond to FTY720 treatment

Anastellin is known to block sphingosine-1-phosphate receptor 1 (S1PR1) signaling (37). This is of particular interest since S1P1R is a current therapeutic target for the treatment of patients with MS through the clinically useful S1P signaling modulator fingolimod (FTY720) (38, 39). Furthermore, S1P1R expression on astrocytes has been shown to be required for the clinical effectiveness of FTY720 in EAE (40). Knowing anastellin is an S1PR1 antagonist and its abundance is increased in the secretome of *Timp1*^cKO^ astrocytes *in vitro*, we wanted to test whether *Timp1*^cKO^ mice with MOG_35-55_ EAE exhibit a different response to FTY720 treatment. MOG_35-55_ EAE WT and *Timp1*^cKO^ mice were administered daily intraperitoneal injections of FTY720 starting at peak clinical illness (18 dpi, blue arrow). We observed that FTY720-treated EAE-WT mice exhibited an accelerated rate of recovery while *Timp1*^cKO^ mice did not exhibit any significant change in clinical profile (**Fig. 4*A*, 4*B***). When compared with genotype-matched untreated EAE groups, the effect of FTY720 in WT mice was significantly different from cKO and treated cKO groups, whereas FTY720-treated *Timp1*^cKO^ mice did not differ from their untreated counterparts (**Fig 4*B***). To be certain that the clinical responses in WT and *Timp1*^cKO^ mice were not due to differences in the phosphorylation of FTY720 (41), and since astrocytes are a key CNS cell type mediating this, we applied FTY720 to primary astrocytes in culture, measured both FTY720 and its metabolite (active form) *phospho-(p)FTY720* in the media by HPLC. We found that the time course of FTY720 metabolism to pFTY720 was equivalent in both WT and *Timp1*^cKO^ astrocytes (**Fig 4C**), therefore, the absence of TIMP1 did not impact the active metabolism of FTY720.

**Figure 4.**
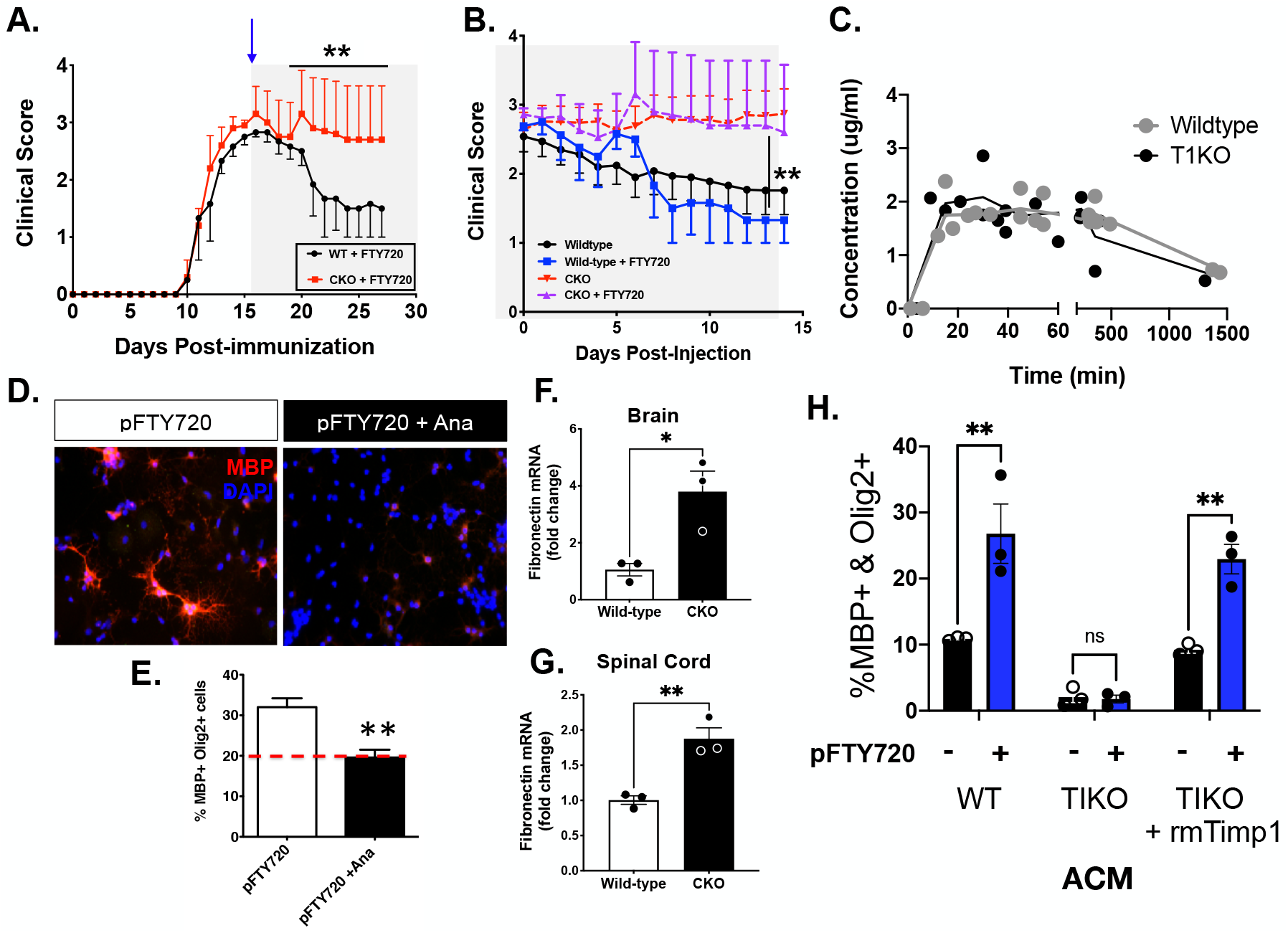
Astrocyte TIMP-1 ckO mice exhibit elevated Fn production and attenuated responses to FTY720. **(A)** EAE clinical scores of Wt and GFAP-Cre:TIMP-1^fl/fl^ mice with FTY720 treatment (initiated by shaded region on graph and blue arrow) demonstrate the lack of clinical response in TIMP-1cKO mice. **(B)** Comparative analysis of FTY720 effect plotted as differential in clinical scores relative days post-injection. (Data in A,B Repeated measures ANOVA, P<0.0001; where post-hoc **, P<0.03-0.006 d21-28). **(C)** Comparative analysis of FTY720 metabolism in Wt and GFAP-Cre:TIMP-1^fl/fl^ primary astrocytes cultures determined differences in astrocytic metabolism of FTY720 compound between cells of either genotype. **(D,E**) Analysis of OL differentiation in primary OPC cultures treated with pFTY720 or pFTY720 and rmAnastellin. Dashed line in E represents baseline level of OL differentiation under control conditions. Data in E was analyzed by t-test relative to baseline differentiation to determine significance **, P<0.008. **(F, G**) Analysis of Fn mRNA expression in brain (t-test, *, P<0.05) and and spinal cord (t-test, **, P<0.01) tissues from EAE Wt and GFAP-Cre:TIMP-1^fl/fl^ mice at time of peak clinical illness (Day 18). **(H**) Analysis of OL differentiation following treatment of pFTY720 treated OPCs with ACM (WT, T1KO, T1KO+rmTimp1). Data in H was analyzed by a 2-way ANOVA. Graphs represent data from n=3-6 technical replicates from an experiment which was representative of n=3 biologically replicated experiments.

We have previously shown that *Timp1*^KO^ EAE mice develop a chronic demyelinating phenotype (23) and since S1PR1 signaling is also known to directly stimulate OPC differentiation (42), we reasoned that elevated expression of Fn in this context could contribute to the block in oligodendrocyte differentiation in these mice as well. To test whether Anastellin alone could act as an inhibitor of S1PR1 signaling on OPCs, we applied pFTY720 (1 nM) onto rOPC cultures under differentiating conditions either with or without coincident treatment with rmAna (1 μg). Quantification of mature OLs (MBP+/Olig2+) revealed that application of pFTY720 alone increased the number of mature MBP+/Olig2+ OLs whereas simultaneous treatment with rmAna completely blocked this effect (**Fig. 4*D*, 4*E***). These data suggest that Anastellin is a novel Fn peptide that can regulate rOPC differentiation and may contribute to the influence of astrocytes lacking TIMP1 production on OPC differentiation and CNS remyelination *in vivo*. Taken together, these data point to a role for astrocyte derived TIMP1 in the regulation of Fn and oligodendrocyte differentiation potential.

We next wanted to determine whether fibronectin (Fn) expression was also differentially affected in the CNS of *Timp1*^cKO^ mice during EAE. We found higher levels of *Fn* expression in both the brains and spinal cords of *Timp1*^cKO^ mice when compared to WT littermates (**Fig 4*F*, 4*G***). These data suggest that astrocyte-specific deletion of *Timp1* altered fibronectin in the CNS during EAE and supported *Timp1*^cKO^ astrocytes as a source of dysregulated fibronectin expression. To verify that it was TIMP-1 that was responsible for the regulation of Fn, and by extension Ana production, which influenced FTY720 responses, we returned to the controlled conditions of primary astrocytes in culture where we tested whether ACM from rmTIMP1 treated *Timp1*^cKO^ astrocytes could rescue the ability of the *Timp1*^cKO^ ACM to promote rOPC differentiation in the presence of pFTY720. To test this, we collected ACM from *Timp1*^cKO^ astrocytes treated with rmTIMP1 (10 ng/mL) or control (veh), and added it to OPCs treated with pFTY720 (1 ng) or control (veh). The percentage of differentiated (MBP+/Olig2+) rOPCs present in groups treated with the *Timp1*^cKO^ + rmTIMP1 ACM was significantly higher than the OPCs treated with only *Timp1*^cKO^ ACM, and was not significantly different from WT (**Fig. 4H**). In the pFTY720 treated rOPCs, the percentage of rOPC differentiation was overall increased in the WT and *Timp1*^cKO^ + rmTIMP1 ACM treatment groups and remained significantly higher than the *Timp1*^cKO^ ACM treatment group (**Fig. 4H**). Overall, these findings show that supplementing the *Timp1*^cKO^ ACM with rnTIMP1 in the extracellular environment can restore the beneficial influences that astrocytes have on promoting rOPC differentiation.

### Production Imbalance between Fn and TIMP-1 in Multiple Sclerosis astrocytes

Lastly, to determine whether the observed changes in fibronectin and TIMP1 in murine astrocytes and CNS tissues were consistent with expression in human MS, we analyzed *TIMP1* and *FN1* mRNA expression in primary astrocytes from human post-mortem MS autopsy specimens (26). Analysis of the ratio of *FN1/TIMP1* expression in these astrocyte cultures showed no spontaneous differences in expression either at the mRNA (**Fig 5*A***) or protein expression levels (**Fig 5*B***). We then tested whether application of recombinant human TIMP1 (rhTIMP1) affected the expression of FN in these human astrocytes. We found that rhTIMP1 significantly reduced the level of FN expression in MS but not control astrocytes (**Fig 5*C***). These data suggest that human astrocytes from MS patients are responsive to TIMP1 and expression of FN in astrocytes from MS disease may have a more labile FN expression pattern that is sensitive to TIMP1 levels.

**Figure 5.**
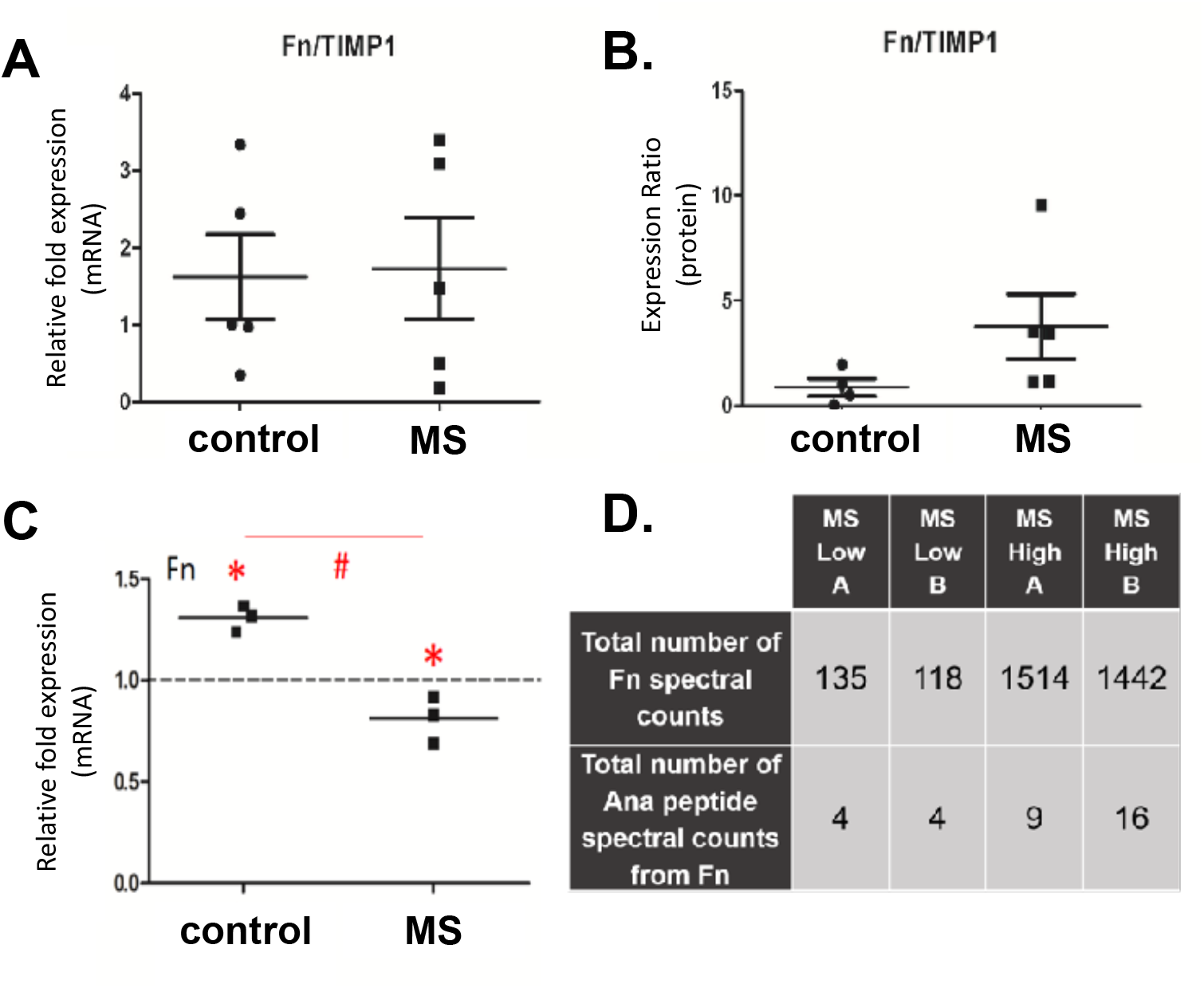
Characterization of Fn and TIMP-1 expression in primary human astrocytes and identification of Anastellin in human MS proteomic databases. **(A)** Ratio of Fn mRNA to TIMP-1 mRNA expression in normal versus MS brain tissue as analyzed by qRT-PCR. **(B)** Ratio of Fn protein to TIMP-1 protein as analyzed by western blot. **(C)** Expression of Fn protein in normal human and MS astrocytes after TIMP-1 treatment. Dotted line indicates no treatment (control). Red asterisks indicate significant difference from control (p<0.05). Pound sign indicates a significant difference between the two groups (p<0.05). **(D)** Proteomic analysis reveals Fn and Ana in the cerebral spinal fluid (CSF) of MS patients [10]. “Low” indicates low cytokine levels in the CSF, while “High” indicates high cytokine levels in the CSF. A and B represent technical replicates.

Since there are no available reagents to specifically and uniquely measure the production of Anastellin itself without also recognizing FN, we examined available proteomics databases for FN spectral counts and the Anastellin peptides among MS patient samples. This analysis confirmed previous studies which identified increased FN in MS, but also found evidence for elevated Anastellin peptides among MS patients with high disease activity scores (**Fig 5*D***), providing a potential association between Anastellin and disease activity in MS.

## Discussion

In this study we found that loss of astrocytic TIMP-1 expression results in changes to the astrocyte secretome that is characterized by elevated abundance of the fibronectin and the fibronectin-derived peptide, Anastellin. We demonstrate that Anastellin can directly impair OPC differentiation and attenuate the action of FTY720 to promote OPC differentiation. We also find that cell-specific deletion of TIMP-1 from astrocytes results in elevated CNS levels of fibronectin and a worsened clinical EAE phenotype that is also unresponsive to FTY720 treatment. These data establish a novel association between TIMP-1 regulation of astrocytes, fibronectin and an astrocyte influence on clinical EAE. Importantly, this was observed in the human MS setting demonstrating that increased Anastellin tracked with disease incidence.

Previous studies have identified TIMP-1 as a protein produced by activated astrocytes (15, 19, 31). TIMP-1 expression during development of the CNS is higher than in the adult CNS where it is only expressed at very low to undetectable levels of expression. In contrast, elevated astrocytic expression of TIMP-1 has been associated with several neurological diseases, including MS (6, 12, 21, 43), HIV-1 encephalitis (44), ischemic brain injury (45, 46), and aging and Alzheimer’s disease (47). The objective of this study was to understand how astrocytic expression of *Timp1* influences these astrocyte responses and how the lack of TIMP-1 production can contribute to CNS pathology. To explore the contribution of TIMP-1 from astrocytes we generated a TIMP-1 transgenic mouse by which we could selectively ablate the TIMP-1 allele in a cell-specific manner. This approach could therefore advance our understanding of TIMP-1 in a range of disease models, from Alzheimer’s to virus infections. In this report we find that the conditional loss of TIMP-1 expression from astrocytes results in a worsened clinical disease course in an EAE model. Using proteomics, we determined that a consequence of astrocytic TIMP-1 loss is dysregulated post-translational processing of fibronectin. We found that fibronectin was a contributing factor to the failure of TIMP-1 knockout astrocytes to promote OPC differentiation. Moreover, our PF-2D proteomic approach enabled us to identify that is was not just fibronectin, but a differentially generated fibronectin peptide product called Anastellin that likely underlies the lack of clinical response to FTY720 treatment in GFAP-Cre:TIMP-1^fl/fl^ mice.

Anastellin was first identified as a peptide derived from the first type III repeat of fibronectin and was named for its ability to inhibit tumor cell growth and angiogenesis of xenografted human cancer cell lines (35). Subsequent study determined that the Anastellin peptide can inhibit sphingosine-1 phosphate (37). These findings prompted us to examine more closely the association of Anastellin as a biologically active counterpart of fibronectin related to the knockout TIMP-1 and its effects on astrocytes and CNS myelination. One impact of this study is the determination that Anastellin, likely through its actions as an endogenous modulator of S1PR1 signaling, could effectively nullify the therapeutic effects of FTY720 (Fingolimod) *in vitro* and in a mouse model of inflammatory demyelination. Hypothetically, based on the results herein, translation of these findings would suggest that assaying Anastellin levels directly could provide an *a priori* means of determining whether an individual may benefit therapeutically from FTY720 treatment. However, it is worth noting that study of Anastellin is made difficult by the lack of specific reagents to differentially identify it from its parent molecule, Fibronectin. It is for this reason that direct measurement of Ana is possible currently only by MS/MS analysis, but the biological effects of Ana have been instead studied by the application of recombinant Ana peptide to biological systems. Hence, development of Anastellin-specific reagents would be a necessary tool to test the translational potential of Ana as a putative biomarker in this and other systems in which its actions have been identified. Moreover, based on our findings, future studies will be needed to elucidate the cell type(s) Anastellin is acting upon during EAE. Current data would indicate that Anastellin may act upon astrocytic S1PR1 is critical for the therapeutic actions of FTY720 (40). Our *in vitro* data support this concept but also demonstrated that Anastellin can act as a direct inhibitor of OL differentiation. Hence, the extracellular production of Anastellin resulting from loss of astrocytic TIMP-1 would be expected to have promiscuous effects on multiple cell types in which S1P signaling has been implicated.

The findings of this study support emerging and increasingly important role for astrocytes and fibronectin in multiple sclerosis (26). Previous work by others has shown that Anastellin promoted high molecular weight (HMW) aggregations of fibronectin (36). Our identification of Anastellin provides a compelling association between the presence of Anastellin in MS we have identified and the HMW fibronectin aggregates found within MS lesions that are thought to impair remyelination (26). When considered in the context of astrocytic TIMP-1, we now hypothesize that TIMP-1 influences the proteolysis of fibronectin, and its loss of which leads to fibronectin breakdown products such as Anastellin, which may then promote the formation of HMW aggregates reported within MS lesions (26) (**Fig. 6**). The mechanism responsible generating Anastellin at this time is unclear. Given the well characterized function of TIMP-1 as a metalloproteinase (MMP) inhibitor, one would surmise that elevated MMP activities would result from the loss of astrocytic TIMP-1. In our prior study (23), we did not find this to be the case, and more recently, elevated MMP-7 activities are associated with clearance of fibronectin aggregates (48), hence TIMP-1 regulation of MMP-related proteolysis does not fully explain our findings. Nevertheless, our work confirms that TIMP-1 impacts the inhibitory effect of fibronectin on OPCs and this is enhanced by anastellin-like peptide(s). Thus, reduced expression of TIMP-1 in conditions such as MS may precipitate important changes to the cellular environment which link reduced remyelination potential and therapeutic clinical responsiveness to astrocytes in this disease.

**Figure 6.**
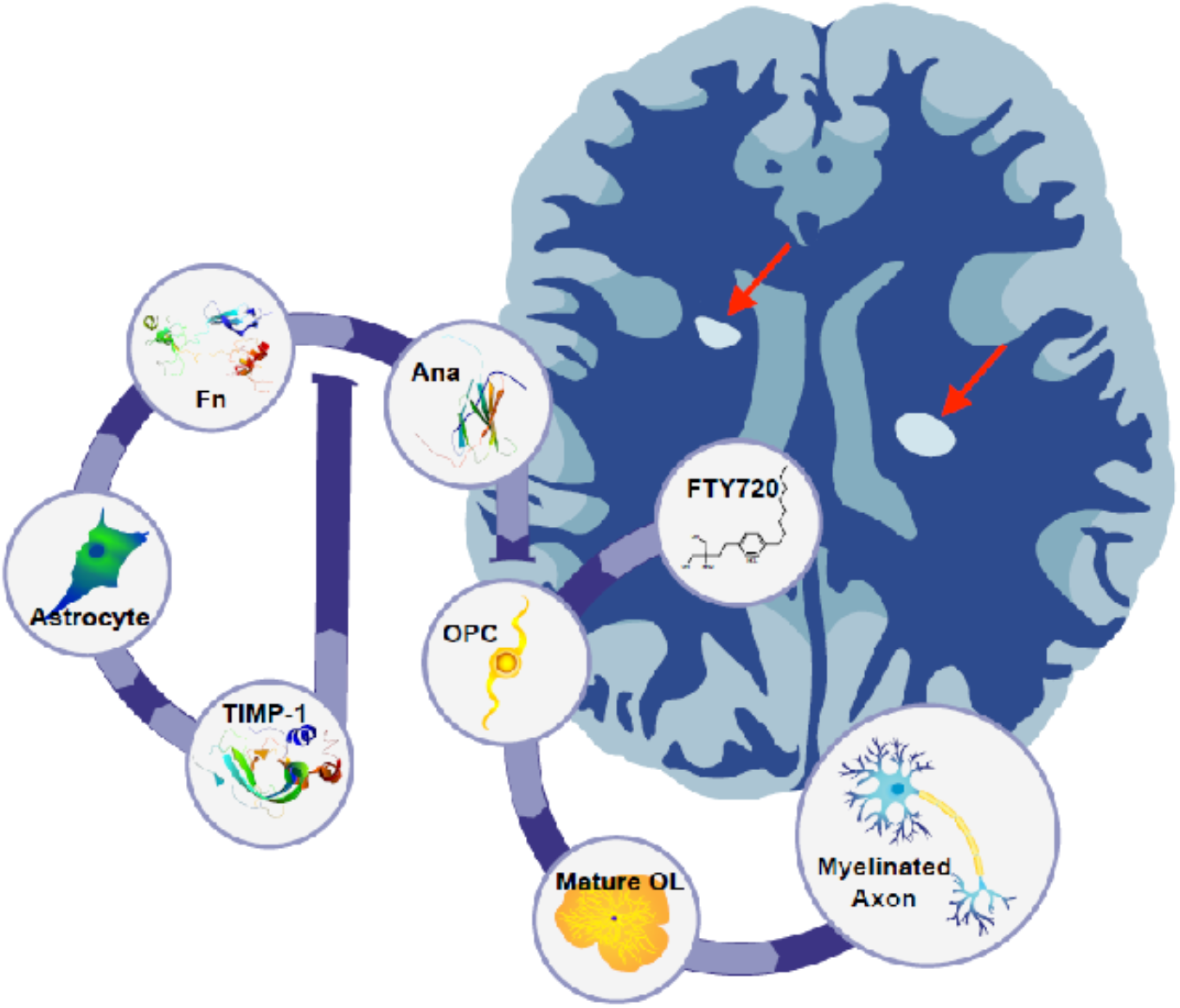
Proposed model for astrocytic TIMP-1 regulation of oligodendrocyte progenitor cell differentiation by Anastellin. Astrocytes when stimulated express and release TIMP-1, which under non-disease conditions would limit the production of Anastellin from fibronectin. In the absence of astrocytic TIMP-1 the production of Anastellin impairs and maintains OPCs differentiation and blocks the ability of FTY720 to promote differentiation. Together, these factors culminate in reduced remyelination of axons contributing to chronic demyelinating plaques in the CNS of MS patients.

Failure of remyelination is now regarded as an underlying cause of clinical disease progression in MS because chronic demyelination contributes to neurodegeneration and progressive clinical disability (49). Our data indicate that a paucity of TIMP-1 expression in MS may be an indicator of inadequate remyelinating potential and failure to express TIMP-1 may therefore be a contributing factor to progression in MS (14). Central to this hypothesis that loss of astrocytic TIMP-1 contributes to chronic demyelination in MS is the question of why expression of TIMP-1, which is the typically robustly produced by astrocytes, would be diminished with chronic inflammation. Prior studies have identified that TIMP-1 expression by astrocytes is not a ubiquitous response to stimulation, but is selective to factors such as serum and interleukin-1β (31), and the induction of TIMP-1 is lost with sustained stimulation by these inflammatory signals (50). More recently, oncostatin M has been shown to also induce astrocytic TIMP-1 and it is TIMP-1 that mediated the regenerative effects of this IL-6 family cytokine on CNS remyelination (21). The present study builds upon these prior studies by corroborating astrocytes as a critical source of TIMP-1. These data are also consistent with prior studies which correlate the induction of TIMP-1 with remyelinating potential in humans: in patients with acute demyelinating encephalomyelitis (ADEM) (51), an acute demyelinating condition that resolves with remyelination, wherein elevated levels of TIMP-1 are observed, but in contrast, among patients with MS, a chronic demyelinating condition, levels of TIMP-1 are not elevated above levels measured in healthy, non-disease control subjects (52, 53). Whether elevated Anastellin peptides contribute to vascular changes in MS requires further study. We may now extend the correlation of this TIMP-1 paradigm directly to include the counter-regulation of fibronectin and Anastellin as a reflection of the human disease condition. Indeed, our finding that Anastellin is generated by astrocytes lacking TIMP-1 may reflect a chronic stage of disease is also consistent with the known function of Anastellin as an angiogenesis inhibitor (35), and reduced angiogenesis observed in chronically demyelinated lesions in MS (54). Taken together, these data suggest that astrocytes likely contribute importantly to disease progression and therapeutic options.

## Materials and Methods

### TIMP-1 Conditional Knockout Mice

A novel TIMP-1 conditional knockout targeting vector containing LoxP sites flanking exons 2 through 5 was generated and injected into mouse embryonic stem cells to generate chimeric mice. The mice were bred with ROSA26-FLPe mice to remove the neomycin cassette. The resulting TIMP-1 fl/fl mice were crossbred with mice expressing Cre recombinase in glial fibrillary acidic protein positive (GFAP+) cells (mGFAP-Cre transgenic line 77.6 [JAX Stock #012887 B6.Cg-Tg(Gfap-cre)77.6Mvs/J]), creating astrocyte-specific TIMP-1 conditional knockout mice. Mice are genotyped to confirm the deletion.

### Primary Mixed Glial Cultures

Animal protocols were performed in compliance with the University of Connecticut Health Center Institutional Animal Care and Use Committees (IACUC). Primary mixed glial cultures were prepared, as previously described. Briefly, cortices of mouse pups (postnatal day 0-3) were obtained from wild-type C57BL/6 mice or TIMP-1 knockout (TIMP-1 KO) mice of the same genetic background, stripped of meninges, separated from the cerebellum, and hippocampi removed. The cortices were dissociated using a neural tissue dissociation kit according to the manufacturer’s protocol (Miltenyi Biotec, Auburn, CA) and cultured in Dulbecco’s Modified Eagle Medium (DMEM) culture medium (Gibco/Life Technologies, Carlsbad, CA). Media containing non-adherent cells was removed the following day and replaced with fresh media. For experiments analyzing the effects of reintroducing TIMP-1 protein to TIMP-1 KO cultures, recombinant murine TIMP-1 (rmTIMP-1, R&D Systems, Minneapolis, MN; 10ng/mL) was added 24 hours prior to harvesting of protein/RNA.

### Isolation of Rat Oligodendrocyte Progenitor Cells (OPCs)

Rat OPC cultures were generated, as previously described (55, 56). Cortices were obtained from rat pups (postnatal day 0-2), stripped of meninges, and had hippocampi removed. Cortices were mechanically minced, then manually homogenized with a flame polished pasteur pipet. The cellular suspension was spun at 1800 rpm for 9 min at room temperature and the pellet was resuspended and plated on poly-l-lysine (0.3mg/mL; Sigma Aldrich) coated flasks with DMEM/F12 mixed cell culture medium (F12 media, Gibco). When a confluent monolayer of cells was formed, flasks were placed in an incubated shaker at 55 rpm for one hour. Media was changed to fresh F12 media to remove cell debris and microglia. Cultures were allowed to sit in a 37°C incubator for 2-4 hours, followed by shaking at 255 rpm overnight to lift OPCs. During shaking of flasks, glass coverslips were washed in sterile 1N HCl for 30 min at room temperature, followed by coating with poly-l-orithine (0.05mg/mL; Sigma) for 3 hours. Supernatant from shaken flasks was spun down at 1800 rpm for 10 min and pellet was resuspended in F12 media and incubated for 3-4 hours. OPCs contained in F12 media were plated on poly-l-orithine coated coverslips and cultured in either differentiation media (get the contents) or Fn depleted astrocyte conditioned media (ACM) for 4 days; method of depletion is explained below. For cultures that were treated with recombinant murine Anastellin (R&D Systems) and/or recombinant Fibronectin (Sigma-Aldrich) were applied as described using a range of concentrations (μg/ml).

### Human Astrocyte Cultures

Adult human astrocytes were isolated from post-mortem subcortical white matter of healthy subjects and patients with multiple sclerosis, as described previously (57). Astrocyte cultures used were at least 97% pure. The cells were plated in 10 cm dishes (1.0 × 10^6^/dish) or 8-well chamber slides (10,000/well).

### Immunodepletion of Fibronectin

Mixed glial cultures were grown as described above. ACM was collected in a 15mL conical tube, and antisera against Fn was applied to the media (1:100; ab23750, Abcam) and then incubated at room temperature for 30 min to functionally deplete media of Fn. Following depletion, Fn was added to OPC cultures to analyze the effect of Fn on OPC differentiation. A species matched IgG control (normal rabbit IgG; Santa Cruz Biotechnology, Dallas, TX) was added to media, obtained from the same mixed glial cultures, to serve as a control for the immunodepletion.

### Protein-Fractionation Two Dimensional (PF-2D) Proteomic Analysis

Confluent monolayers of wild-type and TIMP-1 KO primary mixed glial cultures were grown for 24 hours in media free of serum and collected. The media was dialyzed in varying concentrations of phosphate buffers to remove the salt from the media. The conditioned media was then run through a size filtration column to separate out the proteins by their molecular weight and collected. Eluted proteins were run through protein fractionation 2-dimensional (PF2D), where the proteins were then further separated by a gradient of isoelectric point and then hydrophobicity, as previously described (Beckman Coulter, Pasadena, CA) (34, 58). Identities of proteins differentially produced in TIMP-1 KO and wild-type mixed glial cultures were determined by mass spectrometry (MS/MS).

### Western Blotting

Western blot analysis was performed, as previously described (22). Briefly, cells from mixed glial cultures were collected and protein was digested using a lysis buffer containing Triton X-100 (Sigma Aldrich, St. Louis, MO) and supplemented with protease and phosphatase inhibitors (Roche Diagnostics, Indianapolis, IN). A PierceTM bicinchoninic acid (BCA) assay (Thermo Scientific, Waltham, MA) was used to determine protein concentration. Protein was loaded onto a precast Mini-PROTEAN TGX SDS polyacrylamide gel (4-15%; Bio-Rad, Hercules, CA) and electrophoresed at 200v for 30 minutes. Protein was transferred to a PVDF membrane by electrophoresis for 2 hours at 100v. Membrane was blocked in a blotting buffer containing NaCl, Tris-HCl, and Tween-20 (Sigma) supplemented with 5% non-fat dry milk for 1 hour. Membrane was exposed to primary antisera in blotting buffer supplemented with 2% milk overnight at 4°C. The primary antibody used to identify Fn protein levels was ab6584 (antirabbit, Abcam). Following wash with blotting buffer, membrane was exposed to secondary antibody against rabbit, conjugated with horseradish peroxidase (1:10,000; Vector Laboratories, Burlingame, CA), in blotting buffer for 1 hour. Following wash, membrane was exposed to Amersham ECL Prime Western Blotting Detection Reagent (GE Healthcare Life Sciences, Pittsburgh, PA) and developed onto CL-XPosureTM UV film (Thermo Scientific). Relative band density of protein was determined using ImageJ Software (NIH) (Tan and Ng, 2008). Fn protein levels were compared to a beta-actin (loading control).

### Immunocytochemistry

OPCs plated on poly-l-orithine coated OPCs, after 4 days of differentiation, were fixed with 4% paraformaldehyde (Sigma) at room temperature for 20 minutes. Cells were blocked with PBS containing 0.3% Tween X-100 (PBS-T) and normal goat serum (5%, Sigma) for 1 hour at room temperature. Cells were then incubated overnight at 4°C with PBS-T containing 2% NGS and antisera against Olig-2 (1:500,Millipore) A2B5 (which stains for OPCs) (1:200, Invitrogen), or MBP (which stains for myelinating oligodendrocytes) (1:500, Millipore). The following day the cells were exposed to Alexa-fluorophore-conjugated secondary antisera (1:500; Invitrogen). and counterstained DAPI (Sigma) for 1 hour at room temperature. Coverslips were mounted with Fluoromount G (Southern Biotech, Birmingham, AL) and visualized using an inverted Olympus IX71 microscope outfitted with digital image capture software.

### Antibodies

Anti-Fibronectin antibody (Biotin) (ab6584) (Abcam, Cambridge, UK; rabbit, 1:5000), Anti-Fibronectin antibody (ab23750) (Abcam; rabbit, 1:100-immunodepletion), Anti-Myelin Basic Protein antibody [12] (MBP) (Millipore; rat monoclonal, 1:500), Mouse anti-A2B5 (Invitrogen, Carlsbad, CA; 3ug/mL), 4’,6-diamidino-2-phenylindole (DAPI) (Invitrogen, 1:1000), Mouse anti-glial fibrillary acidic protein (GFAP) (Millipore, 1:500), Beta-actin (Sigma Aldrich, St. Louis, MO; mouse, 1:10,000). All immunostaining was visualized using Alexa-fluorophore-conjugated secondary antisera (1:500; Invitrogen).

### Quantitative Real-Time Polymerase Chain Reaction (qRT-PCR)

RNA was harvested from primary mixed glial cultures using TRIzol reagent according to manufacturer’s protocol (Invitrogen). Following DNase treatment using a TURBO DNase kit (Life Technologies, Carlsbad, CA), RNA was reverse transcribed into complementary DNA (cDNA) using an iScript cDNA synthesis kit (Bio-Rad). Both procedures were performed according to the manufacturer’s protocol. The PCR reaction was carried out, according to manufacturer’s protocol (Bio-Rad), using Sso FastTMEvaGreen^®^ Supermix. The reaction included primer sequences specific to Fn (Integrated DNA Technologies-IDT, Coralville, IA), constructed according to a previous experiment (forward 5’-AGA CCA TAC CTG CCG AAT GTA G-3, reverse 5-GAG AGC TTC CTG TCC TGT AGA G-3) (Morita et al., 2007). Primers for the housekeeping gene GAPDH were used as an internal control for RNA loading, and used to determine the relative gene expression of Fn, in each RNA sample (forward 5’-ACC ACC ATG GAG AAG GC-3’, reverse 5’-GGC ATG GAC TGT GGT CAT GA-3’) (IDT). qRT-PCR was run using a CFX Connect Real-Time PCR Detection System (Bio-Rad). The comparative cycle threshold analysis (△△CT) was used to determine the relative mRNA expression as previously described (Schmittgen and Livark, 2001).

### Experimental Autoimmune Encephalomyelitis (EAE)

EAE was induced in Wt C57BL/6 and TIMP-1 conditional knockout (TIMP-1 CKO) mice between 8 to 10 weeks of age. Mice were immunized with myelin oligodendrocyte glycoprotein peptide (MOG_35-55_, 3mg/mL). MOG_35-55_ peptide was emulsified in complete Freund’s adjuvant (CFA) containing *Mycobacterium tuberculosis* (4mg/mL). 100μL of the emulsification was injected subcutaneously in the thigh region of each hind leg. Animals received an intraperitoneal injection of pertussis toxin (500ng) at the time of immunization and 2 days following. Mice were evaluated daily to monitor weight change and clinical EAE score. The scoring system is as follows: 0, no signs of ailment; 1, limp tail; 2, mild or unilateral hind limb paresis; 3, full hind limb paralysis; 4, moribund; 5, death due to EAE. Mice were monitored for 52 days following immunization and sacrificed for analysis.

### FTY720 Administration

Wt and TIMP-1 CKO mice received daily FTY720 (3mg/kg) injections beginning at the peak of clinical illness and continued daily for 14 days consecutive days. Peak illness was defined as maintaining a constant score of 2 or above for at least 2 days. Control groups were injected with and equivalent volume of vehicle (phosphate buffer saline, PBS).

### *In vitro* FTY720 Metabolism Analyses

Analysis of FTY720 metabolism in primary astrocyte cultures was performed using flow injection mass spectroscopy based on a previously method by Nirogi, et. al (Bioanalysis, 2017 9(7) 565-577). Briefly, samples (5 uL) were introduced into the mass spectrometer (ABSciex 4000 QTrap LC-MS/MS) using a water/methanol/formic acid solution (50/50/0.1%) at an infusion rate of 250 uL/min. Analytes were detected using positive ESI mode through multiple reaction monitoring (MRM) using the parent/daughter ratios (Q1/Q3) of 388.193/290.100 (FTY720P) and 308.300/255.200 (FTY720). The ion spray capillary voltage was 4000 V and source temperature was set at 650^0^C. Nitrogen was used as the curtain gas (10.00) and the collision-induced dissociation gas in Q2. The ion source gas 1 was set at 23. The declustering potential was 76V, collision energy 15V, and collision exit potential set at 6V. Data was acquired using a dwell time of 150 msec with both mass filters operating in unit mass resolution. Data acquisition and instrument control was carried out using Analyst software and quantification based on peak height external calibration using MultiQuant software.

## Acknowledgements

This work was supported in part by grants from the National Multiple Sclerosis Society (RG-1802-30211 to SJC) and National Institutes of Health (NS78392-2 to SJC). We also acknowledge Maddie Youngstrom and Eugene Miller for expert technical assistance, Hayley Joyal (UConn) for development of the informative summary graphic, TuKiet Lam and Jean Kanyo at the W.M. Keck Foundation Biotechnology Resource Laboratory at Yale University for proteomics services, and Dr. Siu-pok Yee at the Center for Mouse Genome Modification at UConn Health for transgenic mouse development services (*TIMP-1^fl/fl^* mice).

